# Microglial phagocytosis of single dying oligodendrocytes is mediated by CX3CR1 but not MERTK

**DOI:** 10.1101/2023.12.11.570620

**Authors:** Genaro E. Olveda, Maryanne N. Barasa, Robert A. Hill

## Abstract

Oligodendrocyte death is common in aging and neurodegenerative diseases. In these conditions, single dying oligodendrocytes must be efficiently removed to allow remyelination and prevent a feed-forward degenerative cascade. Here we used a single-cell cortical demyelination model combined with longitudinal intravital imaging of dual-labeled transgenic mice to investigate the cellular dynamics underlying how brain resident microglia remove these cellular debris. Following phagocytic engagement, single microglia cleared the targeted oligodendrocyte and its myelin sheaths in one day via a precise, rapid, and stereotyped sequence. Deletion of the fractalkine receptor, CX3CR1, delayed microglia engagement with the cell soma but unexpectedly did not affect the clearance of myelin sheaths. Furthermore, and in contrast to previous reports in other demyelination models, deletion of the phosphatidylserine receptor, MERTK, did not affect oligodendrocyte or myelin sheath clearance. Thus, distinct molecular signals are used to detect, engage, and clear sub-compartments of dying oligodendrocytes to maintain tissue homeostasis.

## INTRODUCTION

Oligodendrocytes produce a lipid-rich multilayered membrane called myelin that aids in rapid and efficient transmission of neuronal action potentials while providing metabolic support to the underlying axon^1,2^. Unlike most cells in the brain, oligodendrocytes are generated throughout life and, if needed, can be replaced via the differentiation of resident oligodendrocyte precursor cells (OPCs)^3–8^. Even with this capacity to be replaced, oligodendrocyte regeneration eventually fails in many neurodegenerative disorders and in early aging^3,9^. Several factors are thought to contribute to this limited repair including a delay or failure to clear cellular and myelin debris^10,11^.

Microglia, the resident phagocytes of the central nervous system, primarily orchestrate the debris clearance process ^12–14^. Microglia rely on many cell surface receptors to detect and respond to dying cells and pathogens in their local microenvironment^15^. Among those, the chemokine receptor CX3CR1, mediates communication between microglia and neurons, allowing for microglia to detect neuronal distress signals by sensing either the membrane bound or soluble form of fractalkine (CX3CL1). The binding of fractalkine is thought to act as a “find me” signal, serving to aid in microglia process migration to the site of injury. Past work has found a role for fractalkine and CX3CR1 signaling in oligodendrocyte development^16–18^ and in models of demyelination^19–21^. How this signaling pathway is involved in the dynamic interaction between individual cells is less clear. Other receptors including tyrosine kinases, such as TYRO3, AXL, and MERTK (TAMs), are responsible for recognizing exposed phosphatidylserine on the outer membrane of apoptotic cells through the aid of binding partners growth arrest-specific 6 (GAS6) and protein S1 (PROS1)^22^. Microglia use this exposed phosphatidylserine as an “eat me” signal, indicating the need for phagocytosis. *Mertk* is expressed in microglia and astrocytes and, like *Cx3cr1*, has been shown to be involved in the microglial response in the cuprizone model of demyelination^23,24^.

While it is well established that microglia are involved in the coordinated phagocytosis of degenerating myelin, how microglia dynamically achieve this at the single cell level in the intact brain is less clear. A combination of *in vivo* cellular manipulation and longitudinal imaging techniques are thus needed^25^. Here we used a new method of single cell demyelination, oligodendrocyte 2-Photon apoptotic targeted ablation (2Phatal)^26–28^, in conjunction with triple and quadruple transgenic mouse lines and longitudinal intravital optical imaging to investigate the dynamic cell-cell interactions and molecular signals involved in the clearance of dying oligodendrocytes and their myelin sheaths.

## RESULTS

### Microglia phagocytose and clear single dying oligodendrocytes in a day

To determine how single dying oligodendrocytes and their myelin sheaths are dynamically cleared from the brain we performed cranial window surgeries on a triple transgenic mouse line that allows for intravital fluorescence imaging of microglia and oligodendrocytes. These mice express a membrane tethered enhanced green fluorescent protein in mature oligodendrocyte and their myelin sheaths (*Cnp-*mEGFP) along with tamoxifen-inducible tdTomato in microglia (*Cx3cr1-*creER:tdTomato) (Figure 1). To cause single oligodendrocyte demyelination we used oligodendrocyte 2Phatal which combines in vivo Hoechst nuclear labeling to photo-bleach the nucleus of a single oligodendrocyte causing DNA damage and subsequent programmed cell death of the targeted cell^26–28^. Importantly, unlike laser ablation, this approach does not cause cell rupture and microglial chemotaxis from the release of intracellular ATP^29^, allowing us to investigate the clearance in the context of programmed cell death^26–28^.

**Figure 1:**
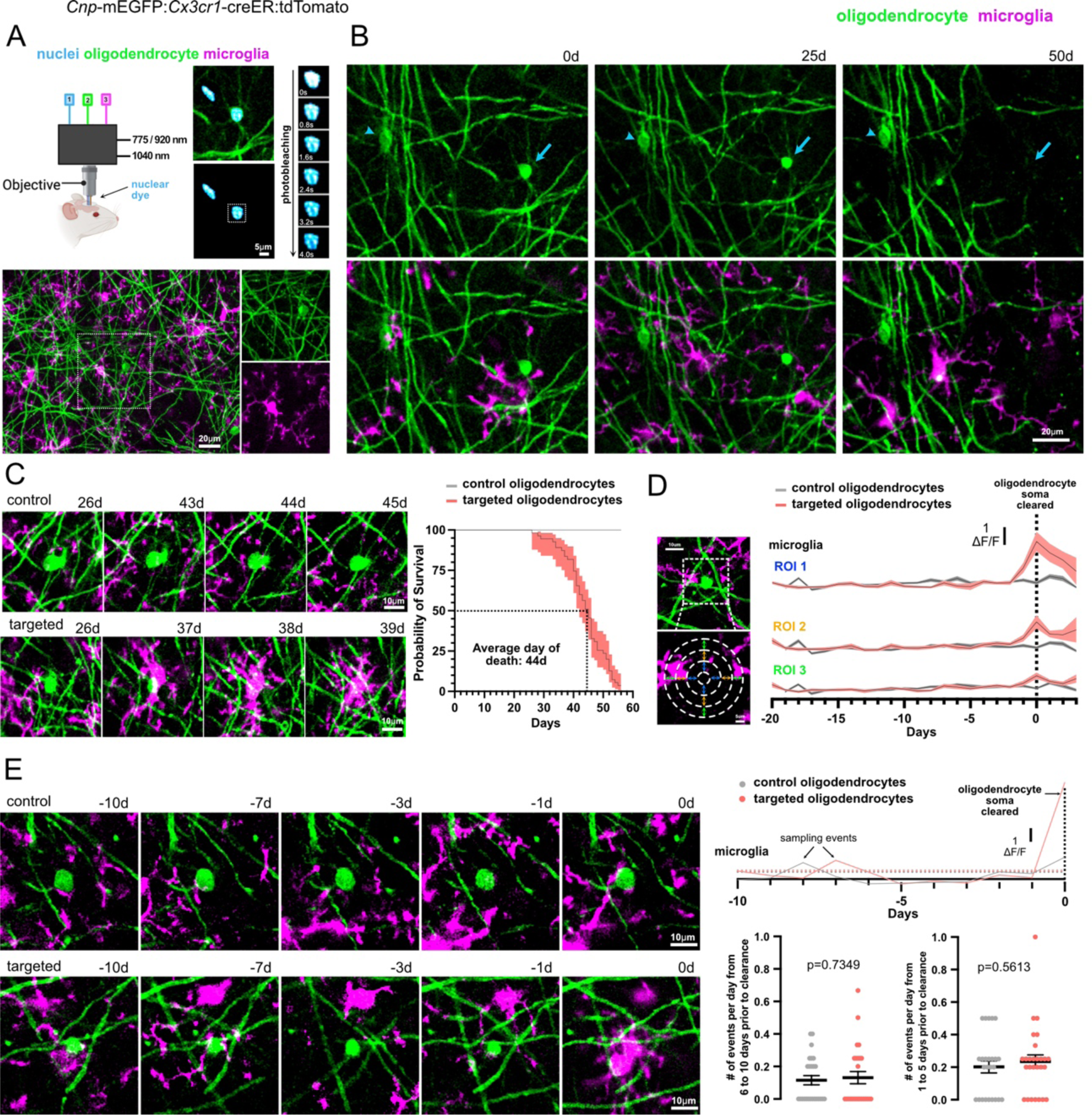
Microglia phagocytose and clear single dying oligodendrocytes in a day. **A)** Single cell ablation of oligodendrocytes, labeled with membrane-tethered EGFP (green) and Hoechst 33342 nuclear dye (cyan), via two-photon apoptotic targeted ablation (2Phatal). Representative image of dual reporter, triple transgenic mice, *Cnp*-mEGFP:*Cx3cr1*-creER:tdTomato. The boxed region shows an individual oligodendrocyte and myelin sheaths (green) and microglia (magenta). **B)** *In vivo* time lapse images showing microglia dynamics surrounding a non-targeted stable oligodendrocyte (arrowhead) and a targeted oligodendrocyte that dies (arrow). **C)** Representative time series of non-targeted (top) and targeted (bottom) oligodendrocytes. Survival curve of non-targeted (gray, n=25) and targeted oligodendrocytes (red, n=56) (the error bands represent the s.e.m). **D)** Analysis of microglia interactions with non-targeted and targeted oligodendrocytes. Microglia fluorescence was measured within 3 regions of interest (ROI) centered around the oligodendrocyte. ROI 1 (3-9um), ROI 2 (9-15um), and ROI 3 (15-21um). Average microglia fluorescence within each ROI 10 days prior to soma clearance and 3 days after. Microglia traces were aligned to the day of soma clearance (day 0). Stable somas were aligned to the average day of soma clearance (day 44). **E)** Time series of microglia interaction events with non-targeted (top) and targeted (bottom) oligodendrocytes. Representative traces of the daily microglia fluorescence from ROI 1 of the non-targeted and targeted oligodendrocytes shown on the left. Sampling events were counted if the microglia fluorescence for that day was one standard deviation (indicated as the dotted line in the graph) above the average microglia fluorescence across all time points for each cell. Quantification of the number of microglia events 6 to 10 and 1 to 5 days prior to soma clearance. (n=25 control cells and 27 targeted cells from 5 mice, unpaired *t*-tests).

Targeted oligodendrocytes died 44 days after single cell DNA damage (Figure 1A-C, Figure S1, n= 56 cells from 5 mice), remarkably consistent with the timing of oligodendrocyte death observed in previous work in other transgenic mouse lines^28^. To directly examine how and when microglia engage, phagocytose, and clear dying cells we imaged mice daily starting at 26 days after 2Phatal and extending until day 56. This allowed us to capture 27 cell soma clearance events. These events were analyzed in depth regarding microglial dynamics leading up to and including the clearance event (Figure 1, Video S1-S2). To determine how microglia dynamically surveilled targeted oligodendrocyte soma we quantified microglia sampling events around control neighboring non-targeted and 2Phatal targeted oligodendrocytes by measuring changes in microglia fluorescence intensity at radiating regions of interest (ROIs) from the center of the oligodendrocyte soma (see methods). These ROIs were defined as: #1 = 3-9µm, #2 = 9-15µm, and #3 =15-21µm (Figure 1D). The microglia fluorescence intensity measurements in each ROI were aligned to the day of oligodendrocyte soma clearance for targeted cells, while those of non-targeted oligodendrocytes were aligned to day 44, the average day of soma clearance for targeted cells. These data revealed that microglia only engaged with the dying oligodendrocytes on the day of clearance and not before (Figure 1D). To formally quantify this, we compared the number of sampling events between 1 to 5 and 6 to 10 days prior to oligodendrocyte soma clearance revealing no significant differences in the number of events between microglia and targeted/non-targeted oligodendrocytes in either of these time windows (Figure 1E, n= 25 non-targeted and 27 targeted cells from 5 mice, unpaired *t*-test). Thus, microglia engage and fully clear single dying oligodendrocytes within a 24-hour window but do not show any bias towards these cells preceding this phagocytic event. Out of the 27 cell soma clearance events that were captured, all of them were carried out by single microglia suggesting that this was not a chemotactic-like response by many microglia but instead followed a programmed cell death mechanism coincident with an orchestrated phagocytic event by single microglia.

### Microglia soma translocation does not impact clearance efficiency

We often observed microglia phagocytosing oligodendrocytes either via **1)** engulfment by microglia processes through a method we called process extension or **2)** engulfment by the microglia soma through a method we called microglia migration (Figure S2). These different methods were apparent in the microglia fluorescence intensity analyses with soma migration events showing brighter signals during the engulfment (Figure S2). However, there were no significant differences the time to oligodendrocyte soma clearance between process extension vs microglia migration (Figure S2B n = 27 process extension events and n = 34 soma migration events from 12 mice). Moreover, there were no differences in the number of microglia sampling events preceding the clearance for these two methods (Figure S2B-C). Therefore, while single microglia used different techniques to engulf the dying oligodendrocyte, there were no overt differences in the clearance timing or efficiency.

### Precise myelin sheath phagocytosis precedes rapid remyelination

We next investigated whether the microglia dynamics involved in clearing a degenerating myelin sheath are like those involved in clearing the oligodendrocyte soma. The clearance of the myelin sheath was observed, on average, 40 days after 2Phatal (Figure 2A-B n= 79 sheaths from 5 mice). To quantify microglia engagement and sampling with control and targeted sheaths, we measured fluorescence intensity changes in microglia in a subset of sheaths at expanding ROIs away from the sheath. These ROIs were defined as: #1 = 0-1.5µm, #2 = 1.5-5µm, #3 = 5-10µm, and #4 = 10-15µm (Figure 2C). In addition to measuring microglia fluorescence within these ROIs, *Cnp*-mEGFP intensity was measured as an indicator of myelin sheath degeneration and reappearance if remyelination occurred. Measurements of 16 degenerating and 18 stable sheaths revealed that microglia engage with and sample both stable and degenerating sheaths (Figure 2D-E). There were no significant differences in microglia engagement with stable or degenerating sheaths preceding the clearance of the sheath (Figure 2E). Notably, microglia engaged only on the day of demyelination exhibiting distinct processes elongation and precise engulfment of the sheath (Figure 2A, D-E, Figure S3-S4, Video S3). This elongation and peak in microglia intensity correlated precisely with the day of demyelination. Like the clearance of the oligodendrocyte soma, microglia efficiently cleared degenerating sheaths within a 24-hour window as evident by the loss of the mEGFP signal (Figure 2D, Figure S3-S4, Video S3).

**Figure 2:**
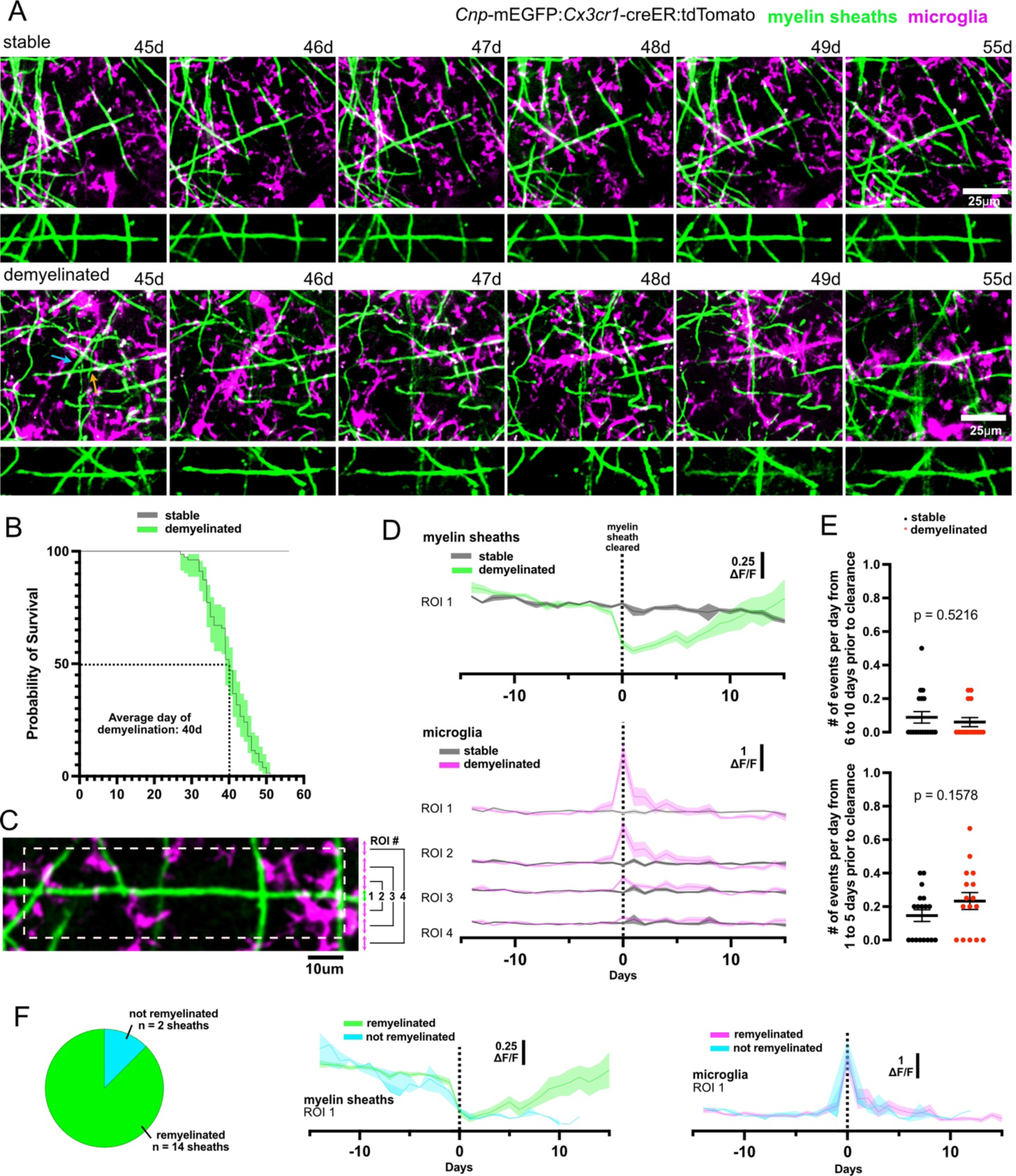
Myelin sheath phagocytosis occurs in a day and precedes remyelination. **A)** In vivo images of a stable sheath (top) and two degenerating sheaths (yellow and blue arrows) that are cleared by single microglia (bottom cropped image shows the sheath indicated by the yellow arrow). **B)** Survival curve of stable (gray, n=18 sheaths from 5 mice) and degenerating sheaths (green, n=79 sheaths from 5 mice) (the error bands represent the s.e.m). **C)** Analysis of microglia interactions with stable and degenerating sheaths. Microglia fluorescence was measured within 3 regions of interest (ROI) centered around the myelin sheath. ROI 1 (1.5um), ROI 2 (1.5-5um), ROI 3 (5-10um), and ROI 4 (10-15um). **D)** Traces of average GFP fluorescence within ROI 1 (top) and average tdTomato fluorescence (bottom) within each ROI, leading up to and following sheath clearance. **E)** Total number of microglia sampling events that occurred within 6 to 10 (top) and 1 to 5 days (bottom) prior to sheath clearance. Measurements were taken from ROI 2. n=18 control sheaths and n=16 targeted sheaths from 5 mice, unpaired *t*-test. **F)** Proportion of degenerated sheaths that were remyelinated and traces of average GFP and tdTomato fluorescence between sheaths that were remyelinated and sheaths that were not remyelinated.

Following the successful clearance of the degenerating sheath, microglia disengaged with the underlying axon, which was followed by remyelination in 88% of the cases shown by the reappearance of an mEGFP labeled myelin sheath (Figure 2F, Figure S3-S4). The dynamics of the microglia engagement was similar between the sheaths that did and did not become remyelinated (Figure 2F). In some cases, remyelination was observed the day after phagocytosis of the sheath (Figure 2A), resembling synchronous remyelination that can occur in young adult animals^28^. Thus, microglia clear degenerating sheaths within a 24-hour period through a coordinated and stereotypical engagement, phagocytosis, and clearance process often resulting in rapid remyelination.

### *Cx3cr1* deletion delays the clearance of single dying oligodendrocytes

*Cx3cr1* is expressed by microglia within the CNS, regulating some aspects of microglia physiology^30–33^ and tissue responses to demyelination^18,20,21^. To determine if CX3CR1 plays in role in the clearance of single dying oligodendrocytes and their myelin sheaths, we performed oligodendrocyte 2Phatal on *Cx3cr1*^+/-^ and *Cx3cr1*^-/-^ mice followed by detailed analysis of soma and sheath clearance.

In *Cx3cr1*^-/-^ mice, targeted oligodendrocytes were cleared on average 48 days after 2Phatal, a 4-day delay compared to *Cx3cr1*^+/-^ mice (Figure 3A-B n = 56 cells from 5 *Cx3cr1*^+/-^ mice and n = 35 cells from 4 *Cx3cr1*^-/-^mice). Moreover, the *Cx3cr1*^-/-^ mice exhibited increased microglia sampling events in the 5 days leading up the clearance (Figure 3C-D, n = 25-27 cells from 5 *Cx3cr1*^+/-^ mice and n = 12 cells from 4 *Cx3cr1*^-/-^mice). This increased sampling was not observed in the 6-to-10-day window suggesting altered microglia response precisely during the 4-day delay when the oligodendrocyte soma would normally be cleared. Despite the significant delay to engage the oligodendrocyte and increased sampling events in *Cx3cr1*^-/-^ mice, after engagement single microglia still cleared the targeted cell within a 24-hour period (Figure 3C-D). Together these data show that deletion of CX3CR1 leads to a delay in the detection of the targeted oligodendrocyte and that during this delay the microglia exhibit increased sampling behavior. After engagement however, the digestion and removal of the cell corpse still happens within a 24-hour period.

**Figure 3:**
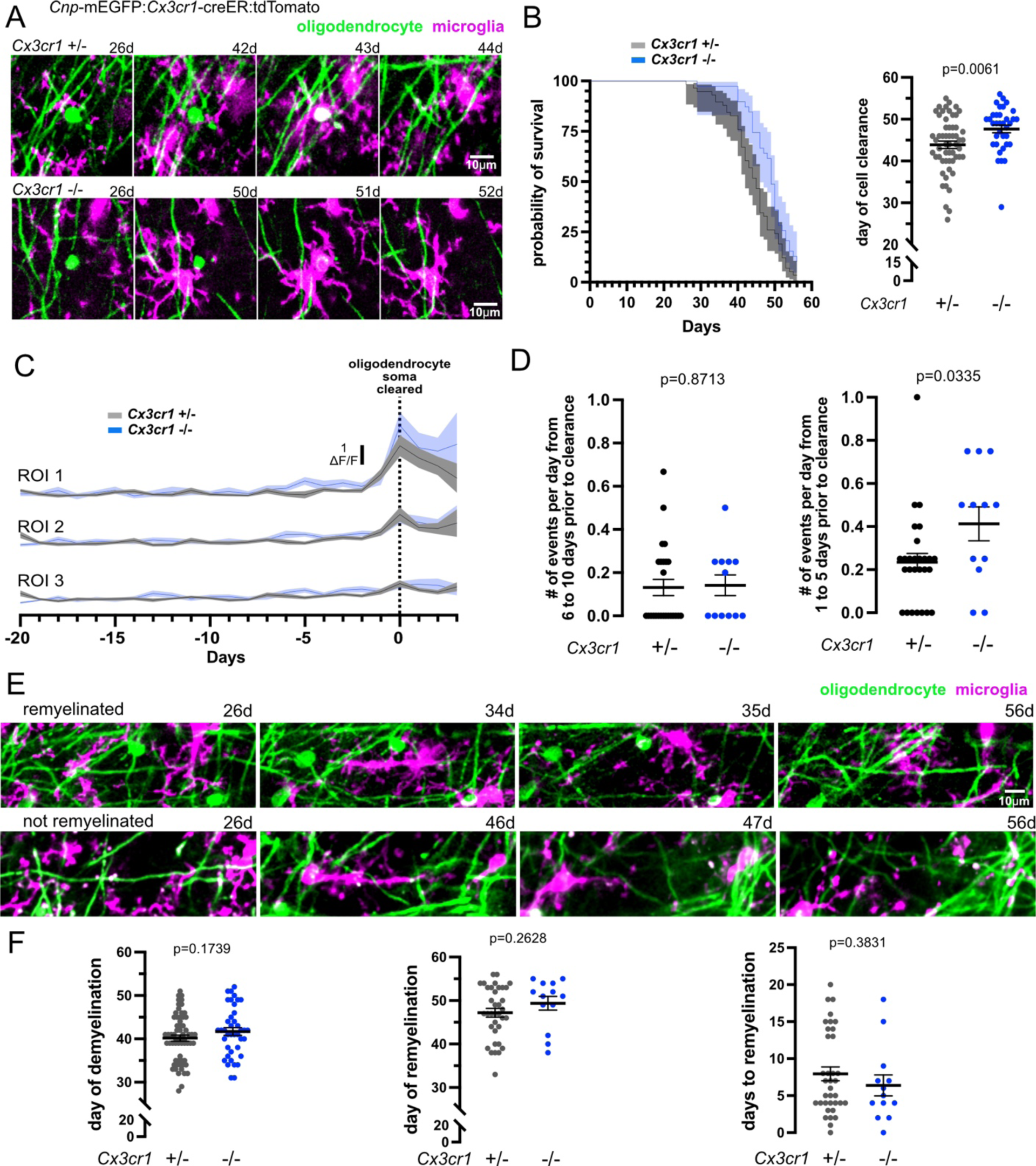
*Cx3cr1* deletion delays the clearance of single dying oligodendrocytes. **A**) Representative time series of oligodendrocyte phagocytosis in *Cx3cr1*^+/-^ and *Cx3cr1*^-/-^ mice. **B)** Survival curve of oligodendrocytes after 2Phatal in *Cx3cr1*^+/-^ (grey) and *Cx3cr1*^-/-^ (blue) mice. Quantification of the average day of soma clearance shows a delay in clearance in the *Cx3cr1*^-/-^ mice, n=56 cells from 5 *Cx3cr1*^+/-^ mice, and n=35 cells from 4 *Cx3cr1*^-/-^ mice, unpaired *t*-test. **C)** Average microglia fluorescence within each ROI 20 days prior to soma clearance and 3 days after in *Cx3cr1*^+/-^ and *Cx3cr1*^-/-^ mice. Microglia traces were aligned to the day of soma clearance (day 0). **D)** The number of sampling events 6 to 10 and 1 to 5 days prior to soma clearance in *Cx3cr1*^+/-^ and *Cx3cr1*^-/-^ mice. n=25-27 cells from 5 *Cx3cr1*^+/-^ mice, and n=12 cells from 4 *Cx3cr1*^-/-^ mice, unpaired *t*-test **E)** Representative time series showing degeneration of a myelin sheath followed by remyelination (top) or failure to remyelinate (bottom). The sheaths that were analyzed were those that intersected a 50μm diameter circle centered around a targeted oligodendrocyte. **F)** Quantification shows no difference in the day of demyelination, day of remyelination, and days to remyelination between *Cx3cr1*^+/-^ and *Cx3cr1*^-/-^ mice, n=63 demyelinated and 36 remyelinated sheaths from 5 *Cx3cr1*^+/-^ mice, and n=40 demyelinated and 13 remyelinated sheaths from 4 *Cx3cr1*^-/-^ mice, unpaired *t*-test.

We next determined whether *Cx3cr1* deletion altered how myelin sheaths were cleared. We investigated microglia dynamics, demyelination timing, remyelination efficiency, and timing to remyelination. First, we analyzed the timing to demyelination within *Cx3cr1*^+/-^ and *Cx3cr1*^-/-^ mice. To do this we determined the fate of myelin sheaths that intersect a 50µm diameter ROI surrounding targeted oligodendrocytes (see methods). A total of 207 sheaths, 115 from *Cx3cr1*^+/-^ and 92 from *Cx3cr1*^-/-^ mice, were selected randomly at day 25 and monitored for stability, demyelination, and potential remyelination by day 56. These sheaths were comprised of a mixed population of those attached to targeted and non-targeted oligodendrocytes, therefore some will remain stable throughout. On average the day of demyelination occurred 40 days post 2Phatal in *Cx3cr1*^+/-^ and 42 days post 2Phatal in *Cx3cr1*^-/-^ mice. Of those sheaths that degenerated, 57% (*Cx3cr1*^+/-^) and 33% (*Cx3cr1*^-/-^) were remyelinated. There were no differences in the average day of remyelination or the time the axon spent demyelinated (Figure 3F n = 63 demyelinated and 36 remyelinated sheaths from 5 *Cx3cr1*^+/-^ mice and n = 40 demyelinated and 13 remyelinated sheaths from 4 *Cx3cr1*^-/-^, mice). Taken together these data show that *Cx3cr1* deletion leads to a significant delay in the clearance of single dying oligodendrocytes, however it has no impact on the timing to detect and clear a degenerating myelin sheath.

### *Mertk* is not required for the clearance of oligodendrocytes or myelin sheaths

*Mertk* has been shown to play a role in clearance of cell corpses and subcellular compartments through phosphatidylserine exposure on the extracellular membrane of dying or damaged cells^22,27^. Moreover, recent work found a role for *Mertk* in microglial responses in the cuprizone model of demyelination which correlated with remyelination efficiency^24^. Therefore, we next determined whether *Mertk* deletion impacted the clearance of single dying oligodendrocytes and myelin sheaths.

Unexpectedly mice lacking the MERTK receptor (*Cx3cr1*^+/-^ *Mertk*^-/-^) showed no delay in the clearance oligodendrocytes compared to *Cx3cr1*^+/-^ *Mertk*^+/+^, with the average being 45 days after 2Phatal (Figure 4B n = 56 cells from 5 *Cx3cr1*^+/-^ *Mertk*^+/+^ mice and n = 31 cells from 2 *Cx3cr1*^+/-^ *Mertk*^-/-^, mice). However, in mice lacking both *Cx3cr1* and *Mertk* (*Cx3cr1*^-/-^ *Mertk*^-/-^) there was a 4-day delay in oligodendrocyte soma clearance, compared to *Cx3cr1*^+/-^ *Mertk*^+/+^ mice (Figure 4B, n = 56 cells from 5 *Cx3cr1*^+/-^ *Mertk*^+/+^ mice and n = 31 cells from 3 *Cx3cr1*^-/-^ *Mertk*^-/-^, mice), remarkably consistent with the delay that was observed in the *Cx3cr1* only knockout mice (Figure 3). We also measured and compared the microglia sampling dynamics across the genotypes and found that all microglia were able to clear the oligodendrocyte soma within a 24-hour window with no difference in the number of sampling events preceding the engagement (Figure 4D n = 25-27 cells from 5 *Cx3cr1*^+/-^ *Mertk*^+/+^ mice, n = 12 cells from 2 *Cx3cr1*^+/-^ *Mertk*^-/-^ mice, and n = 10 cells from 3 *Cx3cr1*^-/-^ *Mertk*^-/-^ mice). Given that past studies have suggested effects of *Cx3cr1* or *Mertk* deletion on microglia migration and/or process motility^24,34–37^ we tested whether the two microglia phagocytic methods (Figure S2) resulted in different oligodendrocyte soma clearance dynamics between the genotypes. The overall incidence of these phagocytic methods was not different between the genotypes and the time to clearance was also not different whether the microglia phagocytosed the oligodendrocyte via soma migration or process extension (Figure S2D).

**Figure 4:**
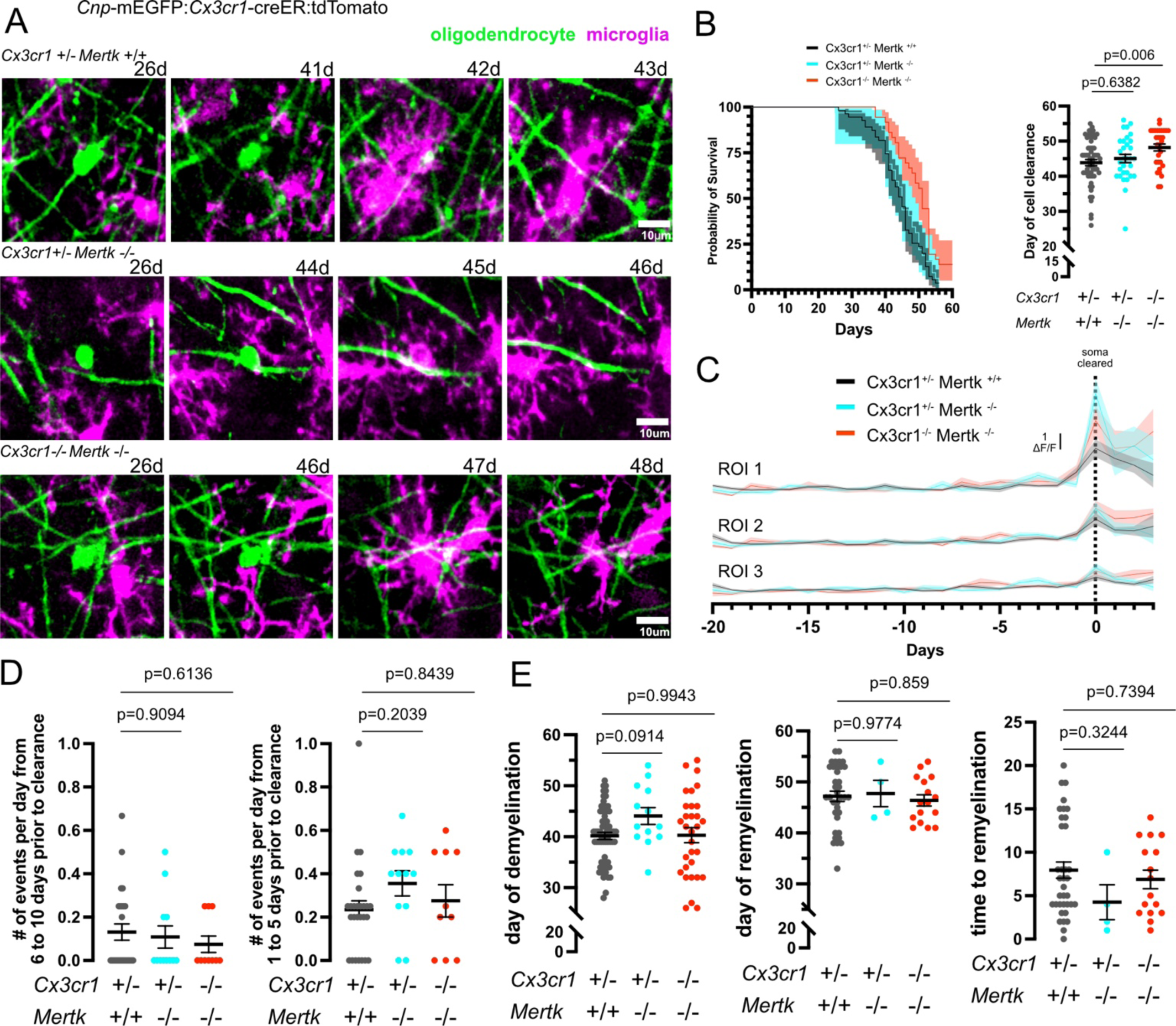
*Mertk* is not required for the clearance of oligodendrocytes or myelin sheaths. **A)** Representative time series of oligodendrocyte phagocytosis in *Cx3cr1*^+/-^ *Mertk*^+/+^, *Cx3cr1*^+/-^ *Mertk*^-/-^, and *Cx3cr1*^-/-^ *Mertk*^-/-^ mice revealing that in all cases the phagocytic event occurs in 1 day. **B)** Survival curve of oligodendrocytes after 2Phatal in *Cx3cr1*^+/-^ *Mertk*^+/+^ (grey), *Cx3cr1*^+/-^ *Mertk*^-/-^ (cyan), and *Cx3cr1*^-/-^ *Mertk*^-/-^ (red) mice. Quantification of the average day of soma clearance shows a delay in clearance in the double knock out mice but not in single *Mertk*^-/-^ mice. (n=56 cells from 5 *Cx3cr1*^+/-^ *Mertk*^+/+^ mice, n=31 cells from 2 *Cx3cr1*^+/-^ *Mertk*^-/-^ mice, and n=31 cells from 3 *Cx3cr1*^-/-^ *Mertk*^-/-^ mice, one-way ANOVA with Dunnett multiple comparisons test. **C)** Traces of the average microglia fluorescence within each ROI 20 days prior to soma clearance and 3 days after. Microglia traces were aligned to the day of soma clearance (day 0). **D)** There were no difference in the total number of sampling events prior to the clearance event across genotypes, n=25-27 cells from 5 *Cx3cr1*^+/-^ *Mertk*^+/+^ mice, n=12 cells from 2 *Cx3cr1*^+/-^ *Mertk*^-/-^ mice, and n=10 cells from 3 *Cx3cr1*^-/-^ *Mertk*^-/-^ mice, one-way ANOVA with Dunnett multiple comparisons test. **E)** Quantification shows no difference in the day of demyelination, day of remyelination, and days to remyelination between the genotypes. The sheaths that were analyzed were those that intersected a 50μm diameter circle centered around a targeted oligodendrocyte. n=63 demyelinated and 36 remyelinated sheaths from 5 *Cx3cr1*^+/-^ *Mertk*^+/+^ mice, n=13 demyelinated and 4 remyelinated sheaths from 2 *Cx3cr1*^+/-^ *Mertk*^-/-^ mice, and n=31 demyelinated and 16 remyelinated sheaths from 3 *Cx3cr1*^-/-^ *Mertk*^-/-^ mice, one-way ANOVA with Dunnett multiple comparisons test.

Similar to no effects in the *Mertk* knockout mice regarding the timing of oligodendrocyte soma clearance, there were also no differences in the day of demyelination, day of remyelination, and time to remyelination (Figure 4E n = 63 demyelinated and 36 remyelinated sheaths from 5 *Cx3cr1*^+/-^ *Mertk*^+/+^ mice, n = 16 demyelinated and 4 remyelinated sheaths from 2 *Cx3cr1*^+/-^ *Mertk*^-/-^ mice, and n = 31 demyelinated and 16 remyelinated sheaths from 3 *Cx3cr1*^-/-^ *Mertk*^-/-^ mice). In the *Cx3cr1*^+/-^ *Mertk*^-/-^ 31% of the sheaths were remyelinated while 52% exhibited remyelination in the *Cx3cr1*^-/-^ *Mertk*^-/-^ mice within the imaging timeframe. Together these data show that *Mertk* expression does not play a significant role in the clearance of single dying oligodendrocytes or their myelin sheaths. These data also replicate the delayed clearance of oligodendrocyte soma when *Cx3cr1* is deleted revealing no additive effect of *Mertk* deletion on the oligodendrocyte soma clearance process.

## DISCUSSION

Here we established a pipeline for the dynamic investigation of how dying oligodendrocytes are cleared in the live mammalian brain. We used this approach to first determine the key checkpoints involved in microglial engagement, phagocytosis, and clearance of both the oligodendrocyte soma and its connected myelin sheaths. Single microglia accomplish this with remarkable precision and speed, in some cases allowing for rapid remyelination to proceed within a day of the phagocytic event. We then went on to investigate (and revisit) the role of two relatively well studied microglial receptors in the clearance of dying oligodendrocytes. Deletion of the CX3CR1 receptor resulted in a delay in clearing the oligodendrocyte soma, but it did not affect the clearance of myelin sheaths. Furthermore, deletion of the MERTK receptor had no impact on microglia dynamics around targeted cells or the clearance of the dying oligodendrocyte soma or myelin sheaths. Collectively, and in the context of past studies, our data illustrate that how, when, and for what reason microglia engage with and clear dying cells and subcellular compartments is highly context and cell-type specific. This has broad impacts on understanding the regulation of tissue homeostasis in normal and pathological contexts.

Past studies that have investigated phagocytosis under demyelinating conditions have primarily used the cuprizone model in fixed tissue. While powerful and well established in the field, this model has several potential limitations, particularly for understanding phagocytosis and debris clearance. First, one should consider what human neurodegenerative condition is being modeled with cuprizone. Cuprizone initiates widespread temporally coordinated intrinsic cell death through likely multiple underlying cell death mechanisms^38–40^. The most prevalent demyelinating disease, multiple sclerosis (MS), is autoimmune mediated, thus widespread intrinsic cell death is not the pathological driver in MS. Cell and myelin debris are certainly generated in MS, but the cellular and inflammatory microenvironment is radically different including blood brain barrier breakdown involving different signaling mechanisms and cell types employed for debris clearance. Indeed, in the most common mouse model of MS, experimental autoimmune encephalomyelitis (EAE), CX3CR1 and TAM receptors have both been shown to impact disease severity. Importantly, however the main reasons for these differences were from disrupted signaling through these receptors on inflammation and activation of peripheral immune cells, not clearance of myelin debris ^41,42^.

Other common neurodegenerative conditions that involve oligodendrocyte death and demyelination include aging and Alzheimer’s disease^43,44,7,45^. Importantly the cell death events in these conditions are not temporally coordinated between many cells, but instead involve protracted death of single cells over weeks to years^7,28,45^. Therefore, the cell debris that must be cleared are not to the scale that occurs with cuprizone, likely engaging different cellular and molecular mechanisms to clear the dying cells and myelin sheaths. Thus, the aging situation is more closely being modeled with 2Phatal^28^ and the lack of significant impacts on myelin sheath phagocytosis and clearance in both the *Cx3cr1* and *Mertk* knockout mice would suggest that these pathways are likely not major contributors to myelin phagocytosis in aging.

Our investigations focused on demyelination and the microglial response in the cortical gray matter. Past work investigating microglial responses to demyelination have focused primarily in white matter areas like the corpus callosum and spinal cord. The distinct neuroanatomical features and cellular compositions between these regions may lead to variations in the microglial response. Thus, it is possible that the disparities between the effects of CX3CR1 and MERTK reported in our study compared to previous studies using the cuprizone model may suggest that the microglia response may be tailored to the specific microenvironment of each region.

The robust and rapid morphological transformation that we observed when single microglia were phagocytosing myelin sheaths resembles past work that has described rod microglia in various neuropathological conditions and aging^46–48^. This past work has pointed out that the function of these rod cells is not entirely clear. Our imaging reveals that this shape transition is surprisingly transient and reflects the shape of the underlying structure that is being phagocytosed by the microglia, in this case the myelin sheath. Thus, the appearance of rod microglia likely reflects the phagocytic events of dendrites, axons, and/or myelin sheaths that are degenerating in situations of damage and/or disease.

While we found no major defects in myelin sheath clearance, we did observe an intriguing delay in the clearance of oligodendrocyte soma when *Cx3cr1* was deleted. This delay was on average 3-4 days, a timeframe that was replicated in the *Cx3cr1 Mertk* double knockout mice. A previous study found a similar ∼3-day delay in the clearance of dying neurons in *Mertk* knockout mice^27^. These findings raise two important concepts. First, the contrast in the receptor needed for the efficient clearance of a cell undergoing programmed cell death from DNA damage, MERTK for neurons and CX3CR1 for oligodendrocytes and second, the similarity in the timing for the “backup” response. In both cases the cell corpse was cleared by microglia, it was merely delayed due to failed initial recognition of the dying cell through phosphatidylserine, in the case of the neuron, or fractalkine, in the case of the oligodendrocyte. One could speculate that there is a 2-3-day window when a cell corpse presents a defined eat me cue before transitioning into another form of cell death and thus presenting an alternate eat me signal that, in the case of the gene knockouts, is now recognized by the microglia resulting in corpse recognition, engulfment, and clearance. The increased number of microglia sampling events in the five days preceding oligodendrocyte soma clearance specifically in the *Cx3cr1* knockout hints that the microglia are sensing the presence of a corpse but are unable to properly engage and phagocytose the cell. While this was not tested in the context of the *Mertk* knockout during neuronal corpse clearance^27^, future work could investigate this hypothesis and further determine the functional consequences of a 3-day delay in corpse clearance to the local tissue inflammation and neural circuitry.

In addition to the cell type dependent differences in phagocytic clearance (neurons vs oligodendrocytes), our data also revealed subcellular differences in the signals regulating the clearance of oligodendrocyte soma vs myelin sheaths. The clearance delay in soma but not myelin sheath observed in the *Cx3cr1* knockout mice suggests that fractalkine may be acting as a signal to aid in the identification of the dying oligodendrocyte soma whereas a different eat me signal may be expressed by the myelin sheath. The source of fractalkine is generally thought to be neurons however recent work suggests the oligodendrocyte lineage also expresses this gene^18^. Both astrocytes and OPCs can also engulf cell structures including myelin^49–51^ and astrocytes and microglia can coordinate to clear different parts of a dying neuron^27^. Given that we were not able to capture all phagocytic engagement events due to their stochastic nature, it is possible that astrocytes and/or OPCs are also involved in the clearance of dying oligodendrocytes, particularly in the context of the receptor knockouts. Future work could use this imaging pipeline and cell death model to explore this question.

Together our data has defined the *in vivo* dynamics involved in the precise microglia dynamics during the clearance of oligodendrocytes and myelin sheaths. Microglia have many receptors that recognize different ligands expressed by dying cells or other cellular debris. The pipeline described here provides a separate framework for investigating this process to further understand when these signaling pathways are used. Future work investigating other receptors including other TAMs, Tyro3^52^ and Axl^53^, TAM binding partners GAS6^54^ and PROS1, and other disease associated receptors such as TREM2^55^, APOE^56^, and SYK^57^ would reveal when and if these pathways are used when individual oligodendrocytes die in the intact brain.

## ACKNOWLEDGEMENTS

We thank members of the Hill lab at Dartmouth for discussions and feedback throughout the project including Tim Chapman and Xhoela Bame for help with preliminary data analysis and imaging. This study was supported by a National Institutes of Health grant no. R01NS122800, a Brain Research Foundation grant no. BRFSG-2019-01, and the Esther A. & Joseph Klingenstein Fund and Simons Foundation to R.A.H. and a National Institutes of Health-NRSA fellowship grant no. F31NS130910 and U.S. Department of Education GAANN fellowship no. P200A210064 to G.E.O.

## AUTHOR CONTRIBUTIONS

G.E.O and R.A.H designed and performed all the experiments, analyzed the data, wrote the manuscript, and secured funding. M.N.B. contributed to image preparation, collation, and quantification. R.A.H supervised the study.

## METHODS

### Animals

All animal procedures were approved by the institutional animal care and use committee at Dartmouth College. The following mouse strains were purchased from Jackson Laboratory and crossed to generate triple and quadruple transgenic mice: *Cnp*-mEGFP^58^ (strain# 026105), *Cx3cr1*-creER^59^ (strain# 020940), *Mertk^-/-^* ^60^(strain# 011122), and floxed tdTomato Ai9^61^ (strain# 007909). For all experiments male and female mice were 3-4 months old at the beginning of the experiment.

### Surgery

For all experiments dual cranial windows were performed, one on each hemisphere. Mice were anesthetized by an intraperitoneal injection of ketamine (100 mg kg^−1^) and xylazine (10 mg kg^−1^). The section of skin above the skull was shaved and sterilized before being removed. A region (3mm x 3mm) of the skull, above the somatosensory cortex, was removed along with the underlying dura. Hoechst 33342 nuclear dye was applied topically (for 2Phatal experiments) to the exposed cortex prior to being sealed with a no. 0 cover glass, allowing for longitudinal *in vivo* imaging. A nut was then glued to the rostral portion of the skull allowing for repeated immobilization for imaging. Mice were given subcutaneous injections of carprofen (50 mg kg^−1^) after surgery, 24hrs, and 48hrs post-surgery.

### Intravital imaging

Mice were anesthetized with isoflurane before being head fixed on the microscope stage. All images were captured using a two-photon microscope (Bruker with Prairieview software) with an InSight X3 femtosecond pulsed laser (Spectra Physics) and a x20 water immersion objective (Zeiss NA 1.0). Lasers were set to following wavelengths to excite Hoechst nuclear dye (775nm), mEGFP (920nm), and tdTomato (1040nm). All image z-stacks were acquired using interlaced scanning with a 303.3µm (X, 1024 pixels), 303.3µm (Y, 1024 pixels), and 52.5µm (Z; step size, 1.5µm) volume. Landmark images were taken of the pial vessels at the surface of the cortex at the beginning of the experiment to relocate the same position over weeks.

### 2Phatal

To induce death of single oligodendrocytes, cranial windows were performed, and Hoechst 33342 was topically applied as stated above. After the surgery, mice were allowed to recover for 24hrs allowing for uniform nuclear labeling. Within each cranial window, three individual positions were identified and imaged using the 775nm, 920nm, and 1040nm laser wavelengths on a two-photon microscope. These channels were then overlayed, allowing for the identification of mature oligodendrocyte nuclei. Single cell death by 2Phatal was performed by placing a single ROI (8µm x 8µm) over the nucleus of a mature oligodendrocyte. Photobleaching of the nuclear dye was done using the 775nm wavelength laser and a dwell time of 100μs. 2phatal was performed in two of the three positions, per window, allowing for an internal control for every mouse. Within the targeted positions, 4 individual *Cnp*-mEGFP labeled mature oligodendrocytes were targeted. Images were acquired before 2Phatal, 25 days after 2Phatal, and every day thereafter for 31 days allowing for targeted oligodendrocyte clearance, demyelination, and remyelination to occur.

### Microglia response to oligodendrocyte death

For all analyses, hyperstack images were created using all images for each position. For cell clearance analysis, cells were marked as cleared at the first time point in which there is no longer any mEGFP signal representing the soma of the targeted oligodendrocyte. To quantify the average day of clearance, targeted oligodendrocytes were monitored from day 25 until the day of soma clearance. Only oligodendrocytes that were cleared within the 56 days were included in the average day of clearance. Non-targeted oligodendrocytes were also monitored throughout this period. To quantify the microglia/oligodendrocyte soma interactions, ROIs with diameters of 3µm, 9µm,15µm, and 21µm, centered around the oligodendrocyte soma in Fiji. Using the “XOR” function, each ROI was combined creating new ROIs, 3-9µm, 9-15µm, and 15-21µm, centered from the oligodendrocyte soma. The “XOR” function combines the selected ROIs by excluding the overlapping area and keeping the non-overlapping regions. Microglia (tdTomato) signal was measured within each of these new ROIs for every timepoint that was captured. Individual microglia traces were aligned to the day of soma clearance. Every timepoint was normalized to the average microglia fluorescence signal for that cell up until the day of clearance. Microglia sampling events were defined when the microglia intensity at any timepoint was one standard deviation from the average intensity for that cell. For these measurements only a single slice from the z-stack was used in which the oligodendrocyte soma was the brightest. Cells were excluded from this analysis if microglia engulfment was not captured the day prior to oligodendrocyte soma clearance.

### Microglia response to myelin sheath degeneration

For the quantification of microglia dynamics surrounding a myelin sheath, both stable and degenerating sheaths were chosen at random in the targeted positions from day 25 images. Stable sheaths were those that were not degenerated throughout the experiment. Similar to the cell death, the myelin sheath was considered degenerated at the first day in which there is no mEGFP signal representing the myelin sheath. For the quantification of changes in myelin integrity de/remyelination, a ROI was created by tracing the myelin sheath using the segmented line tool, 1.5µm width, in Fiji. For every sheath the mEGFP signal was measured within the ROI at every time point. The average trace for the stable sheath was then aligned to the average day of demyelination and the average trace for degenerating sheaths were aligned to the day of degeneration. For the quantification of microglia engagement with stable and degenerating myelin sheaths, the 1.5µm width ROI for every sheath was used as a template to create 5um, 10um, 15um wide ROIs. The “XOR” function in Fiji was used to combine the ROIs together creating new ROIs, 1.5-5µm, 5-10µm, 10-15µm. For every sheath microglia (tdTomato) signal was measured within each ROI and the average traces for the stable and degenerating sheath were aligned to the day of degeneration, respectively. Using the microglia trace from the 1.5-5µm ROI, the number of microglia engagement events were quantified for both stable and degenerating sheaths. Events were calculated the same way as stated previously. This in-depth analysis was only done on *Cx3cr1^+/-^* mice.

For the quantification of the demyelination and remyelination across all genotypes an ROI, 50µm in diameter, was centered around the oligodendrocyte soma. Myelin sheaths that intersect the ROI were selected at the first time point and monitored daily to evaluate the proportion of sheath stability, remyelination, and degeneration without repair occurring within the territory of an oligodendrocyte. Demyelination was classified as the first day of mEGFP lost whereas remyelination was classified as the first day of mEGFP appearance. To quantify the time to remyelination, the time between the disappearance of the myelin sheath to the reappearance of a new myelin sheath was calculated.

### Image Processing

For all analysis, time series images for each position were combined into a hyperstack using Fiji. For oligodendrocyte clearance, individual cells were identified and a single slice, 100µm x 100µm regions was cropped centered around the cell for every time point. These cropped images were then combined into a new hyperstack and aligned using the MultiStackReg plugin in Fiji. Aligned images were then cropped again using a 50um x 50um ROI. For the in-depth sheath analysis individual sheaths were cropped, using a 175µm x 175µm ROI, and a six-optical slice max projection centered at the middle plane of the sheath were made for every timepoint. The stack consisting of max projections were then aligned and cropped using a 100µm x 100µm ROI. The “bleach correction” function in Fiji was then used on the image stack to help restore relative fluorescence level across the image stack. For the demyelination/remyelination analysis, the 100µm x 100µm hyperstacks were used. A 50µm diameter ROI was centered around the oligodendrocyte and only sheaths that intersected this ROI, within two optical slices above and below the middle plane of the soma, were chosen. These sheaths were marked with the multi-point tool in Fiji allowing them to be tracked over time.

### Statistical analyses

All statistical analyses were performed in Prism (GraphPad Software, version 10.1). No statistical methods were used to predetermine sample size. Sample sizes were based on those reported in previous publications. Statistical differences in the number of microglia sampling events (Fig. 1-3) were determined using an unpaired t-test. For quantification of the day of cell clearance and the demyelination/remyelination analysis in *Cx3cr1*^+/-^ and *Cx3cr1*^-/-^ mice (Fig. 3), unpaired t-test was used to determine statistical significance. For quantification of the day of cell clearance, microglia interaction and the demyelination/remyelination analysis in *Cx3cr1^+/-^ Mertk^+/+^, Cx3cr1^+/-^ Mertk^-/-^,* and *Cx3cr1^-/-^ Mertk^-/-^* mice (Fig. 4), one-way analysis of variance with Dunnett’s multiple comparison test. Statistical differences for the average day of soma clearance by process extension and soma migrations and the proportion of each across all genotypes (Fig. S2D) were determined using two-way analysis of variance with Sidak’s multiple comparison test. For each experiment, cell and animal numbers are indicated in the figure legend. Experimenters were blinded to the genotypes of all the mice during data acquisition and analysis.

---

**Figure S1:**
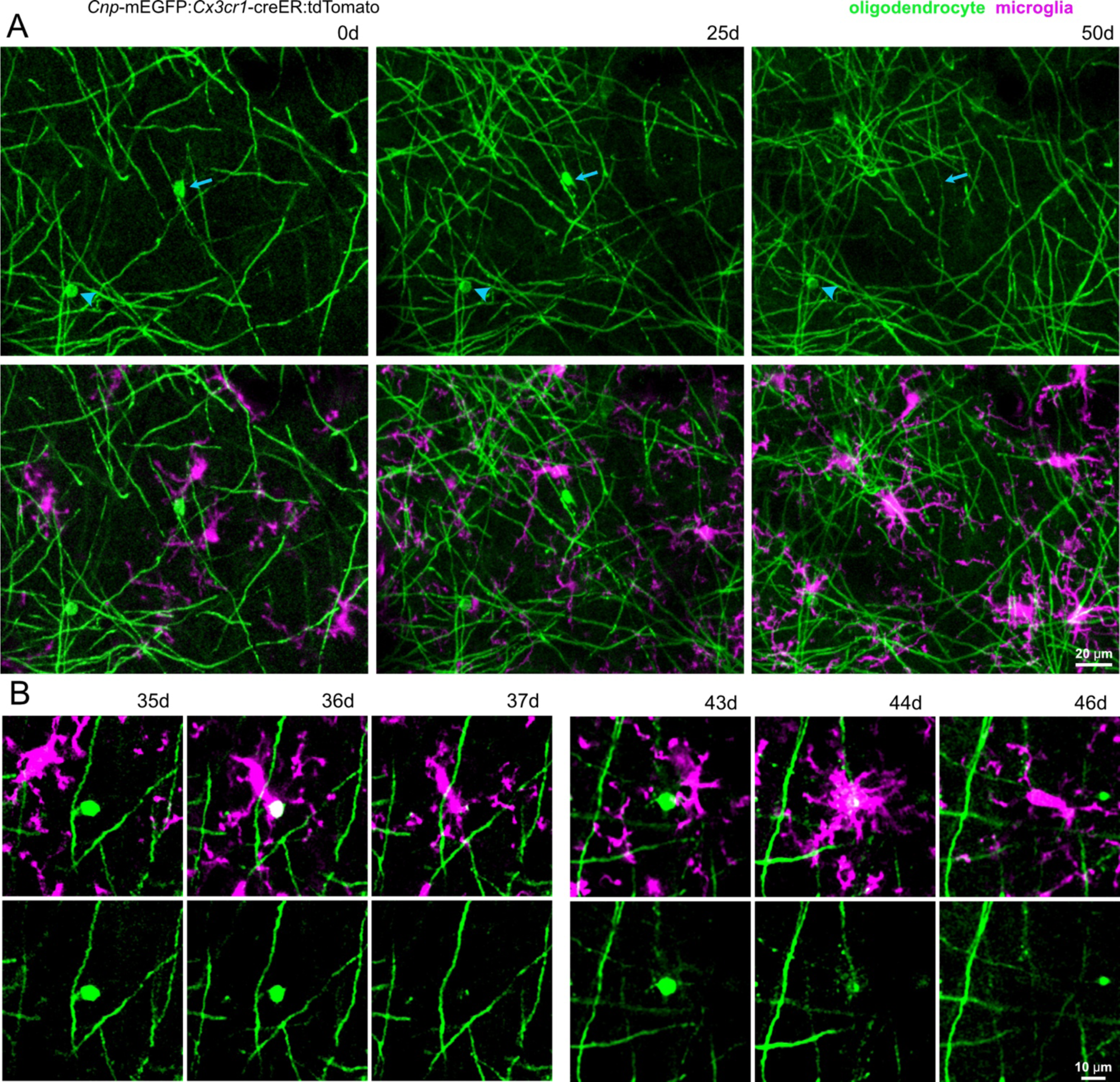
In vivo microglial clearance of single dying oligodendrocytes. **A)** Representative in vivo images from of dual reporter, triple transgenic mice with oligodendrocytes (green) and microglia (magenta) labeling. Example of microglia response to a targeted oligodendrocyte (arrow) and no response to the adjacent non-targeted oligodendrocyte (arrowhead). **B)** Additional representative time series of two separate examples of microglia responding and clearing oligodendrocyte soma within a 24hr window.

**Figure S2:**
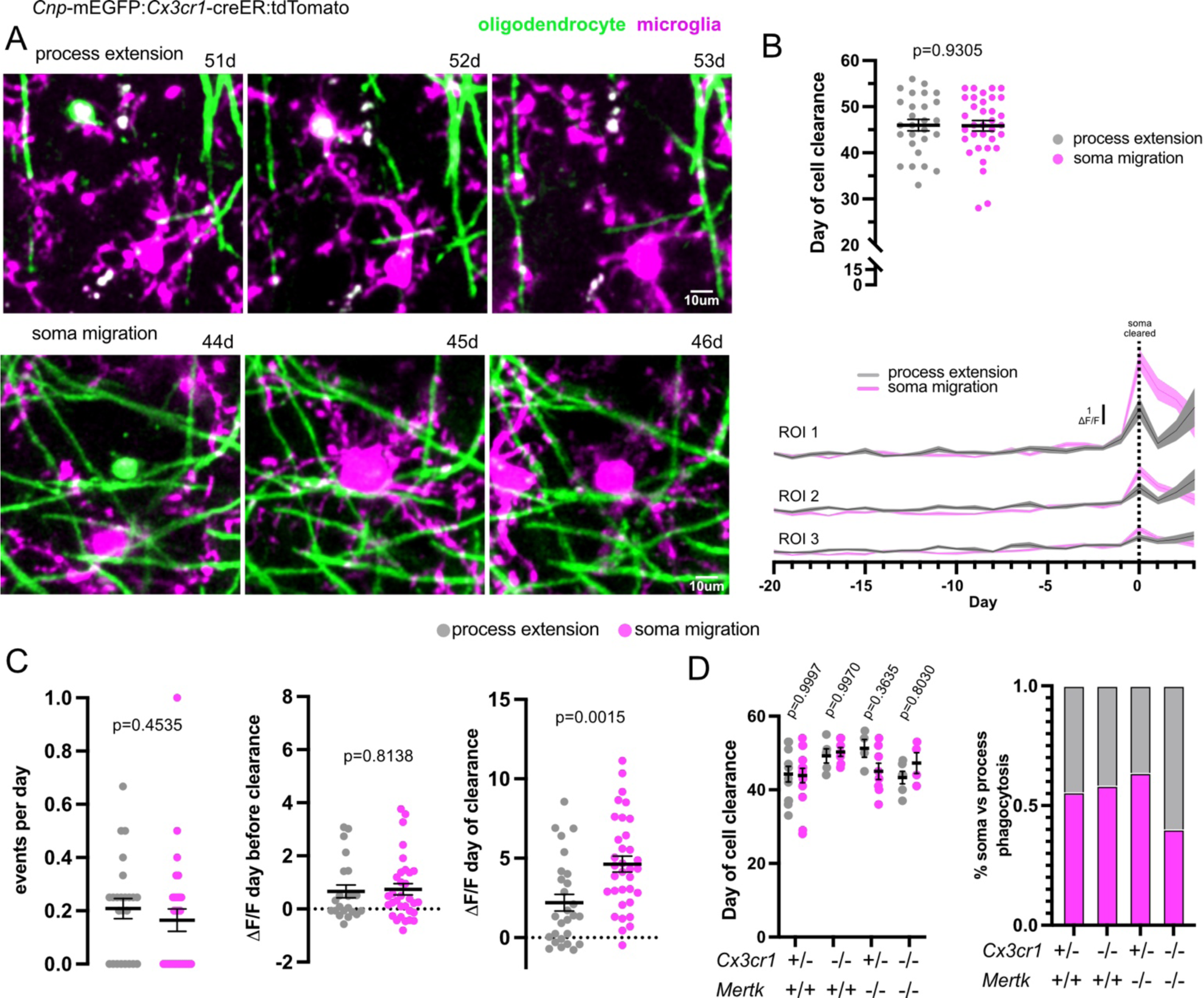
Oligodendrocyte clearance proceeds through two distinct forms of microglia phagocytosis. **A)** Representative time series of oligodendrocyte clearance through microglia process extensions (top) or by microglia soma migration (bottom). **B)** Quantification of the average day of soma clearance shows no difference between clearance via microglia process extension or microglia soma migration. Representative traces of average microglia fluorescence within each ROI 20 days prior to soma clearance and 3 days after. Microglia traces were aligned to the day of soma clearance (day 0). **C)** Quantification of microglia traces in (B). Total number of microglia sampling events that occurred prior to clearance. Microglia fluorescence the day before and day of clearance by microglia process extension (grey) and microglia soma migration (magenta). **D)** Quantification of the average day of soma clearance by process extension and soma migration and the proportion of each across all genotypes, n= 27 cells from 5 *Cx3cr1*^+/-^ *Mertk*^+/+^ mice, n=12 cells from 4 *Cx3cr1*^-/-^ *Mertk*^+/+^ mice, n=12 cells from 2 *Cx3cr1*^+/-^ *Mertk*^-/-^ mice, and n=10 cells from 2 *Cx3cr1*^-/-^ *Mertk*^-/-^ mice.

**Figure S3:**
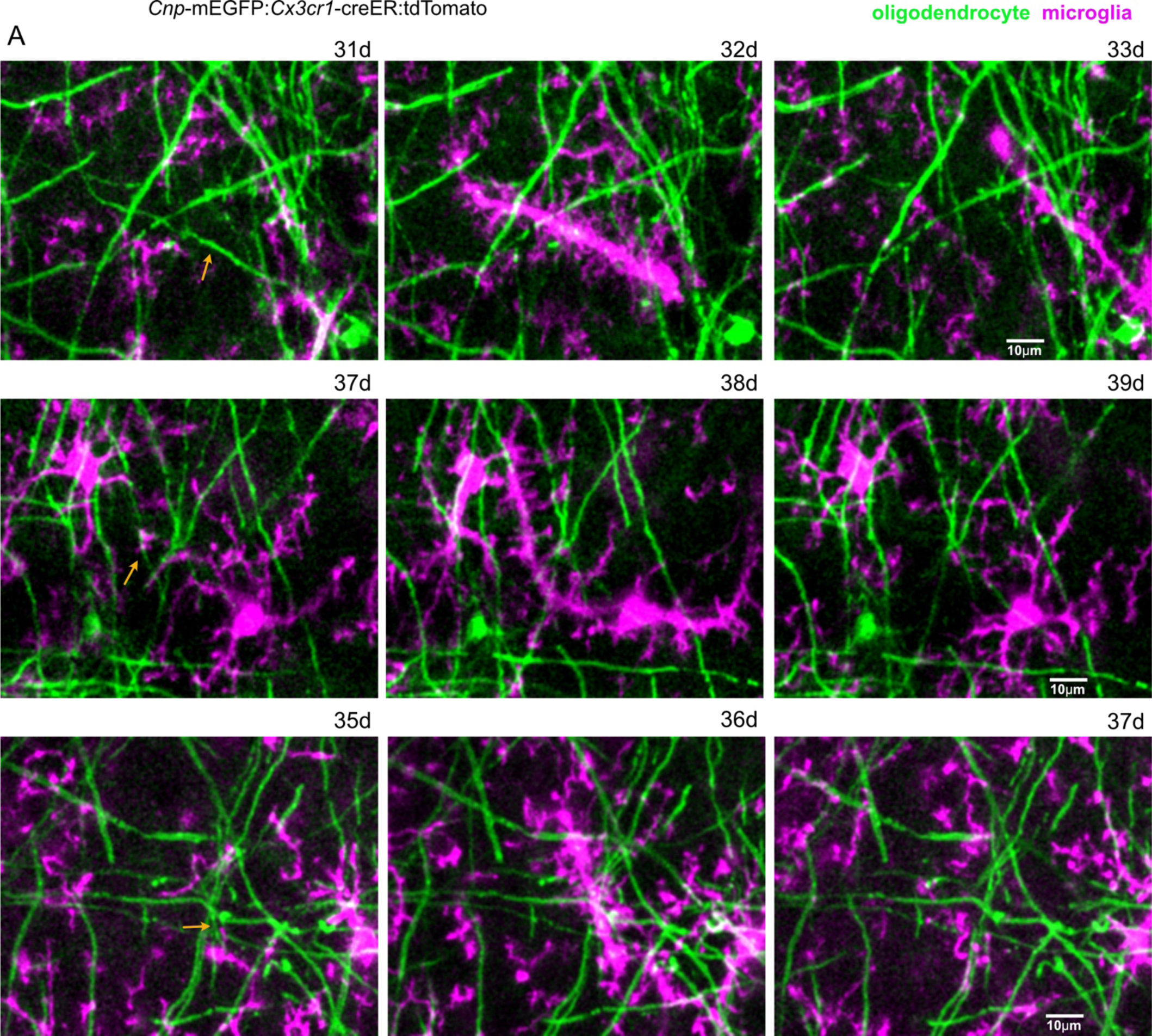
Microglia clear degenerating myelin sheaths in 1 day. **A)** Representative in vivo images showing precise and rapid microglia clearance of demyelinating sheaths (yellow arrows) followed by a rapid disengagement. Myelin sheaths were cleared within a 24hr window.

**Figure S4:**
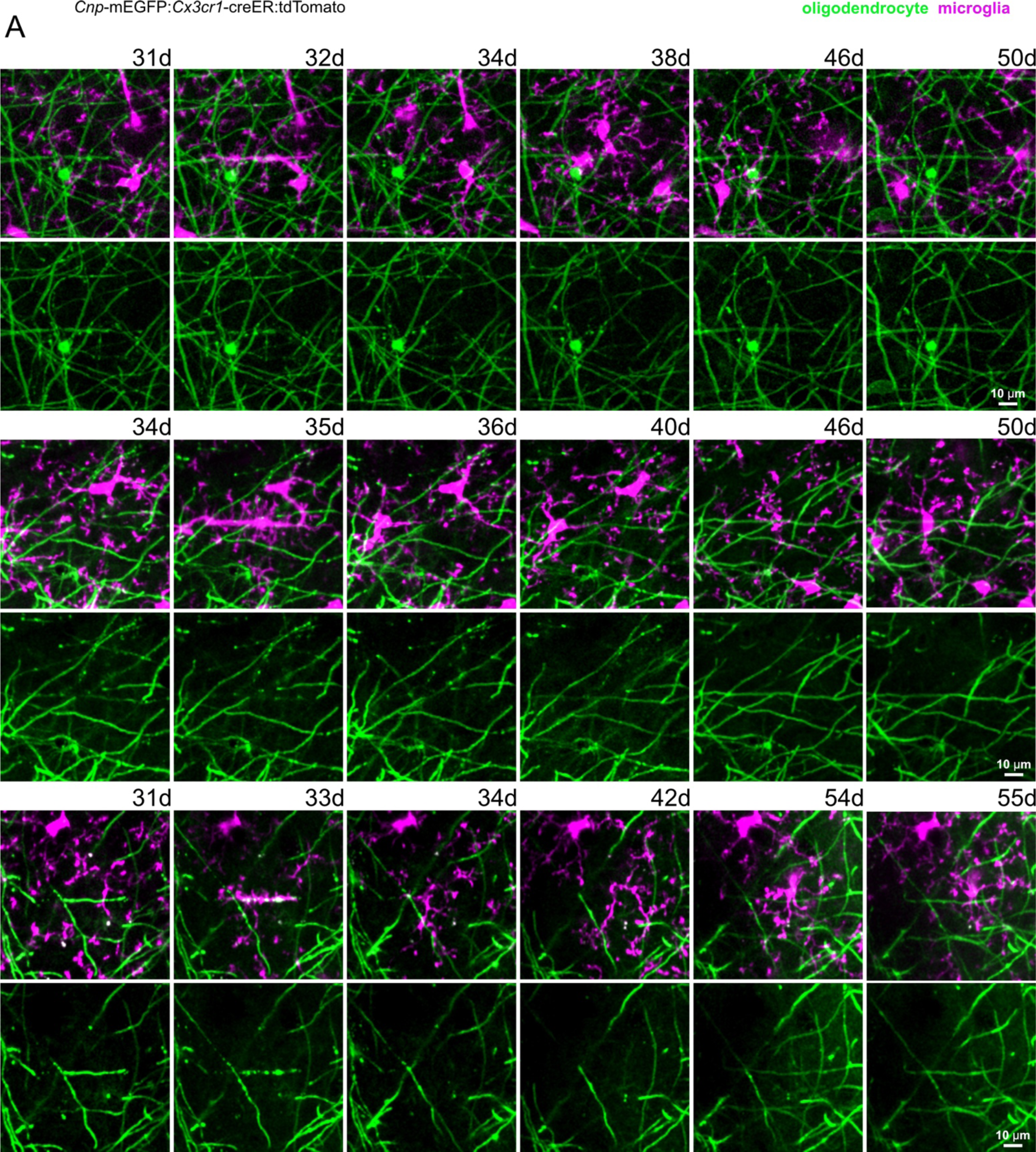
Clearance of myelin sheaths allows for subsequent remyelination. **A)** Additional representative images of sheaths that were degenerated and were remyelinated. The timing of remyelination varied from days to weeks after sheaths were cleared by microglia.

**Video S1: Microglia interactions with oligodendrocyte somas**

In vivo time lapse images of a microglia interacting with a non-targeted (left) and targeted (right) oligodendrocyte. Example of microglia phagocytosis of a dying oligodendrocyte. Video displayed at 3 frames per second.

**Video S2: Microglia phagocytosis of oligodendrocyte somas**

Additional in vivo time lapse images of two adjacent oligodendrocyte somas being phagocytosed by microglia. Video displayed at 3 frames per second.

**Video S3: Microglia phagocytosis of degenerating myelin sheaths followed by remyelination**

In vivo time series showing three examples of microglia phagocytosis of a degenerating myelin sheath. Example one shows microglia clearance of a degenerating sheath (arrow) along with 3 other sheaths that were cleared but microglia engagement was not captured (arrowheads). Note that the mEGFP fluorescence intensity is dim at the initial day of remyelination but increases as the myelin sheath stabilizes. Degenerating sheaths are labeled in white and the remyelination is labeled in gold. Video displayed at 3 frames per second.

## References

1. Salzer, J.L., and Zalc, B. (2016). Myelination. Curr Biol 26, R971–R975. 10.1016/j.cub.2016.07.074.

2. Nave, K.-A. (2010). Myelination and the trophic support of long axons. Nat Rev Neurosci 11, 275–283. 10.1038/nrn2797.

3. Franklin, R.J.M., and Ffrench-Constant, C. (2008). Remyelination in the CNS: from biology to therapy. Nat Rev Neurosci 9, 839–855. 10.1038/nrn2480.

4. Zhu, X., Hill, R.A., Dietrich, D., Komitova, M., Suzuki, R., and Nishiyama, A. (2011). Age-dependent fate and lineage restriction of single NG2 cells. Development 138, 745–753. 10.1242/dev.047951.

5. Kang, S.H., Fukaya, M., Yang, J.K., Rothstein, J.D., and Bergles, D.E. (2010). NG2+ CNS glial progenitors remain committed to the oligodendrocyte lineage in postnatal life and following neurodegeneration. Neuron 68, 668–681. 10.1016/j.neuron.2010.09.009.

6. Young, K.M., Psachoulia, K., Tripathi, R.B., Dunn, S.-J., Cossell, L., Attwell, D., Tohyama, K., and Richardson, W.D. (2013). Oligodendrocyte dynamics in the healthy adult CNS: evidence for myelin remodeling. Neuron 77, 873–885. 10.1016/j.neuron.2013.01.006.

7. Hill, R., Li, A., and Grutzendler, J. (2018). Lifelong cortical myelin plasticity and age-related degeneration in the live mammalian brain. Nat Neurosci 21, 683–695. 10.1038/s41593-018-0120-6.

8. Hughes, E.G., Orthmann-Murphy, J.L., Langseth, A.J., and Bergles, D.E. (2018). Myelin remodeling through experience-dependent oligodendrogenesis in the adult somatosensory cortex. Nat Neurosci 21, 696–706. 10.1038/s41593-018-0121-5.

9. Stadelmann, C., Timmler, S., Barrantes-Freer, A., and Simons, M. (2019). Myelin in the Central Nervous System: Structure, Function, and Pathology. Physiological Reviews 99, 1381–1431. 10.1152/physrev.00031.2018.

10. Kotter, M.R., Li, W.-W., Zhao, C., and Franklin, R.J.M. (2006). Myelin Impairs CNS Remyelination by Inhibiting Oligodendrocyte Precursor Cell Differentiation. J. Neurosci. 26, 328–332. 10.1523/JNEUROSCI.2615-05.2006.

11. Neumann, H., Kotter, M.R., and Franklin, R.J.M. (2009). Debris clearance by microglia: an essential link between degeneration and regeneration. Brain 132, 288–295. 10.1093/brain/awn109.

12. Kettenmann, H., Hanisch, U.-K., Noda, M., and Verkhratsky, A. (2011). Physiology of Microglia. Physiological Reviews 91, 461–553. 10.1152/physrev.00011.2010.

13. Casano, A.M., and Peri, F. (2015). Microglia: multitasking specialists of the brain. Dev Cell 32, 469–477. 10.1016/j.devcel.2015.01.018.

14. Podleśny-Drabiniok, A., Marcora, E., and Goate, A.M. (2020). Microglial Phagocytosis: A Disease-Associated Process Emerging from Alzheimer’s Disease Genetics. Trends in Neurosciences 43, 965–979. 10.1016/j.tins.2020.10.002.

15. Arandjelovic, S., and Ravichandran, K.S. (2015). Phagocytosis of apoptotic cells in homeostasis. Nat Immunol 16, 907–917. 10.1038/ni.3253.

16. Voronova, A., Yuzwa, S.A., Wang, B.S., Zahr, S., Syal, C., Wang, J., Kaplan, D.R., and Miller, F.D. (2017). Migrating Interneurons Secrete Fractalkine to Promote Oligodendrocyte Formation in the Developing Mammalian Brain. Neuron 94, 500–516.e9. 10.1016/j.neuron.2017.04.018.

17. Nemes-Baran, A.D., White, D.R., and DeSilva, T.M. (2020). Fractalkine-Dependent Microglial Pruning of Viable Oligodendrocyte Progenitor Cells Regulates Myelination. Cell Rep 32, 108047. 10.1016/j.celrep.2020.108047.

18. Watson, A.E.S., de Almeida, M.M.A., Dittmann, N.L., Li, Y., Torabi, P., Footz, T., Vetere, G., Galleguillos, D., Sipione, S., Cardona, A.E., et al. (2021). Fractalkine signaling regulates oligodendroglial cell genesis from SVZ precursor cells. Stem Cell Reports 16, 1968–1984. 10.1016/j.stemcr.2021.06.010.

19. Lampron, A., Larochelle, A., Laflamme, N., Préfontaine, P., Plante, M.-M., Sánchez, M.G., Yong, V.W., Stys, P.K., Tremblay, M.-È., and Rivest, S. (2015). Inefficient clearance of myelin debris by microglia impairs remyelinating processes. J. Exp. Med. 212, 481–495. 10.1084/jem.20141656.

20. Mendiola, A.S., Church, K.A., Cardona, S.M., Vanegas, D., Garcia, S.A., Macklin, W., Lira, S.A., Ransohoff, R.M., Kokovay, E., Lin, C.-H.A., et al. (2022). Defective fractalkine-CX3CR1 signaling aggravates neuroinflammation and affects recovery from cuprizone-induced demyelination. J Neurochem 162, 430–443. 10.1111/jnc.15616.

21. de Almeida, M.M.A., Watson, A.E.S., Bibi, S., Dittmann, N.L., Goodkey, K., Sharafodinzadeh, P., Galleguillos, D., Nakhaei-Nejad, M., Kosaraju, J., Steinberg, N., et al. (2023). Fractalkine enhances oligodendrocyte regeneration and remyelination in a demyelination mouse model. Stem Cell Reports 18, 519–533. 10.1016/j.stemcr.2022.12.001.

22. Lemke, G., and Rothlin, C.V. (2008). Immunobiology of the TAM receptors. Nat Rev Immunol 8, 327–336. 10.1038/nri2303.

23. Healy, L.M., Perron, G., Won, S.-Y., Michell-Robinson, M.A., Rezk, A., Ludwin, S.K., Moore, C.S., Hall, J.A., Bar-Or, A., and Antel, J.P. (2016). MerTK Is a Functional Regulator of Myelin Phagocytosis by Human Myeloid Cells. J. Immunol. 196, 3375–3384. 10.4049/jimmunol.1502562.

24. Shen, K., Reichelt, M., Kyauk, R.V., Ngu, H., Shen, Y.-A.A., Foreman, O., Modrusan, Z., Friedman, B.A., Sheng, M., and Yuen, T.J. (2021). Multiple sclerosis risk gene Mertk is required for microglial activation and subsequent remyelination. Cell Rep 34, 108835. 10.1016/j.celrep.2021.108835.

25. Hill, R.A., and Grutzendler, J. (2019). Uncovering the biology of myelin with optical imaging of the live brain. Glia 67, 2008–2019. 10.1002/glia.23635.

26. Hill, R.A., Damisah, E.C., Chen, F., Kwan, A.C., and Grutzendler, J. (2017). Targeted two-photon chemical apoptotic ablation of defined cell types in vivo. Nat Commun 8, 15837. 10.1038/ncomms15837.

27. Damisah, E.C., Hill, R.A., Rai, A., Chen, F., Rothlin, C.V., Ghosh, S., and Grutzendler, J. (2020). Astrocytes and microglia play orchestrated roles and respect phagocytic territories during neuronal corpse removal in vivo. Sci Adv 6, eaba3239. 10.1126/sciadv.aba3239.

28. Chapman, T.W., Olveda, G.E., Bame, X., Pereira, E., and Hill, R.A. (2023). Oligodendrocyte death initiates synchronous remyelination to restore cortical myelin patterns in mice. Nat Neurosci 26, 555–569. 10.1038/s41593-023-01271-1.

29. Davalos, D., Grutzendler, J., Yang, G., Kim, J.V., Zuo, Y., Jung, S., Littman, D.R., Dustin, M.L., and Gan, W.-B. (2005). ATP mediates rapid microglial response to local brain injury in vivo. Nat Neurosci 8, 752–758. 10.1038/nn1472.

30. Bazan, J.F., Bacon, K.B., Hardiman, G., Wang, W., Soo, K., Rossi, D., Greaves, D.R., Zlotnik, A., and Schall, T.J. (1997). A new class of membrane-bound chemokine with a CX3C motif. Nature 385, 640–644. 10.1038/385640a0.

31. Harrison, J.K., Jiang, Y., Chen, S., Xia, Y., Maciejewski, D., McNamara, R.K., Streit, W.J., Salafranca, M.N., Adhikari, S., Thompson, D.A., et al. (1998). Role for neuronally derived fractalkine in mediating interactions between neurons and CX3CR1-expressing microglia. Proc Natl Acad Sci U S A 95, 10896–10901. 10.1073/pnas.95.18.10896.

32. Jung, S., Aliberti, J., Graemmel, P., Sunshine, M.J., Kreutzberg, G.W., Sher, A., and Littman, D.R. (2000). Analysis of fractalkine receptor CX(3)CR1 function by targeted deletion and green fluorescent protein reporter gene insertion. Mol Cell Biol 20, 4106–4114. 10.1128/MCB.20.11.4106-4114.2000.

33. Cardona, A.E., Pioro, E.P., Sasse, M.E., Kostenko, V., Cardona, S.M., Dijkstra, I.M., Huang, D., Kidd, G., Dombrowski, S., Dutta, R., et al. (2006). Control of microglial neurotoxicity by the fractalkine receptor. Nat Neurosci 9, 917–924. 10.1038/nn1715.

34. Maciejewski-Lenoir, D., Chen, S., Feng, L., Maki, R., and Bacon, K.B. (1999). Characterization of fractalkine in rat brain cells: migratory and activation signals for CX3CR-1-expressing microglia. J Immunol 163, 1628–1635.

35. Kj, L., Je, L., Yd, W., W, M., Am, F., Rn, F., and Wt, W. (2009). Regulation of dynamic behavior of retinal microglia by CX3CR1 signaling. Investigative ophthalmology & visual science 50. 10.1167/iovs.08-3357.

36. Pagani, F., Paolicelli, R.C., Murana, E., Cortese, B., Di Angelantonio, S., Zurolo, E., Guiducci, E., Ferreira, T.A., Garofalo, S., Catalano, M., et al. (2015). Defective microglial development in the hippocampus of Cx3cr1 deficient mice. Front Cell Neurosci 9, 111. 10.3389/fncel.2015.00111.

37. Fourgeaud, L., Través, P.G., Tufail, Y., Leal-Bailey, H., Lew, E.D., Burrola, P.G., Callaway, P., Zagórska, A., Rothlin, C.V., Nimmerjahn, A., et al. (2016). TAM receptors regulate multiple features of microglial physiology. Nature 532, 240–244. 10.1038/nature17630.

38. Plemel, J.R., Caprariello, A.V., Keough, M.B., Henry, T.J., Tsutsui, S., Chu, T.H., Schenk, G.J., Klaver, R., Yong, V.W., and Stys, P.K. (2017). Unique spectral signatures of the nucleic acid dye acridine orange can distinguish cell death by apoptosis and necroptosis. J Cell Biol 216, 1163–1181. 10.1083/jcb.201602028.

39. Zirngibl, M., Assinck, P., Sizov, A., Caprariello, A.V., and Plemel, J.R. (2022). Oligodendrocyte death and myelin loss in the cuprizone model: an updated overview of the intrinsic and extrinsic causes of cuprizone demyelination. Mol Neurodegener 17, 34. 10.1186/s13024-022-00538-8.

40. Chapman, T.W., Piedra, E.T., and Hill, R.A. (2023). Oligodendrocyte maturation alters the cell death mechanisms that cause demyelination. Preprint at bioRxiv, 10.1101/2023.09.26.557781 10.1101/2023.09.26.557781.

41. Garcia, J.A., Pino, P.A., Mizutani, M., Cardona, S.M., Charo, I.F., Ransohoff, R.M., Forsthuber, T.G., and Cardona, A.E. (2013). Regulation of adaptive immunity by the fractalkine receptor during autoimmune inflammation. J Immunol 191, 1063–1072. 10.4049/jimmunol.1300040.

42. Gruber, R.C., Ray, A.K., Johndrow, C.T., Guzik, H., Burek, D., de Frutos, P.G., and Shafit-Zagardo, B. (2014). Targeted GAS6 delivery to the CNS protects axons from damage during experimental autoimmune encephalomyelitis. J Neurosci 34, 16320–16335. 10.1523/JNEUROSCI.2449-14.2014.

43. Peters, A., Verderosa, A., and Sethares, C. (2008). The neuroglial population in the primary visual cortex of the aging rhesus monkey. Glia 56, 1151–1161. 10.1002/glia.20686.

44. Bartzokis, G. (2011). Alzheimer’s disease as homeostatic responses to age-related myelin breakdown. Neurobiol Aging 32, 1341–1371. 10.1016/j.neurobiolaging.2009.08.007.

45. Chen, J.-F., Liu, K., Hu, B., Li, R.-R., Xin, W., Chen, H., Wang, F., Chen, L., Li, R.-X., Ren, S.-Y., et al. (2021). Enhancing myelin renewal reverses cognitive dysfunction in a murine model of Alzheimer’s disease. Neuron 109, 2292–2307.e5. 10.1016/j.neuron.2021.05.012.

46. Ziebell, J.M., Taylor, S.E., Cao, T., Harrison, J.L., and Lifshitz, J. (2012). Rod microglia: elongation, alignment, and coupling to form trains across the somatosensory cortex after experimental diffuse brain injury. Journal of Neuroinflammation 9, 247. 10.1186/1742-2094-9-247.

47. Bachstetter, A.D., Ighodaro, E.T., Hassoun, Y., Aldeiri, D., Neltner, J.H., Patel, E., Abner, E.L., and Nelson, P.T. (2017). Rod-shaped microglia morphology is associated with aging in 2 human autopsy series. Neurobiology of Aging 52, 98–105. 10.1016/j.neurobiolaging.2016.12.028.

48. Holloway, O.G., Canty, A.J., King, A.E., and Ziebell, J.M. (2019). Rod microglia and their role in neurological diseases. Seminars in Cell & Developmental Biology 94, 96–103. 10.1016/j.semcdb.2019.02.005.

49. Chung, W.-S., Clarke, L.E., Wang, G.X., Stafford, B.K., Sher, A., Chakraborty, C., Joung, J., Foo, L.C., Thompson, A., Chen, C., et al. (2013). Astrocytes mediate synapse elimination through MEGF10 and MERTK pathways. Nature 504, 394–400. 10.1038/nature12776.

50. Ponath, G., Ramanan, S., Mubarak, M., Housley, W., Lee, S., Sahinkaya, F.R., Vortmeyer, A., Raine, C.S., and Pitt, D. (2017). Myelin phagocytosis by astrocytes after myelin damage promotes lesion pathology. Brain 140, 399–413. 10.1093/brain/aww298.

51. Buchanan, J., Elabbady, L., Collman, F., Jorstad, N.L., Bakken, T.E., Ott, C., Glatzer, J., Bleckert, A.A., Bodor, A.L., Brittain, D., et al. (2022). Oligodendrocyte precursor cells ingest axons in the mouse neocortex. Proceedings of the National Academy of Sciences 119, e2202580119. 10.1073/pnas.2202580119.

52. Blades, F., Aprico, A., Akkermann, R., Ellis, S., Binder, M.D., and Kilpatrick, T.J. (2018). The TAM receptor TYRO3 is a critical regulator of myelin thickness in the central nervous system. Glia. 10.1002/glia.23481.

53. Hoehn, H.J., Kress, Y., Sohn, A., Brosnan, C.F., Bourdon, S., and Shafit-Zagardo, B. (2008). Axl-/-mice have delayed recovery and prolonged axonal damage following cuprizone toxicity. Brain Res 1240, 1–11. 10.1016/j.brainres.2008.08.076.

54. Binder, M.D., Cate, H.S., Prieto, A.L., Kemper, D., Butzkueven, H., Gresle, M.M., Cipriani, T., Jokubaitis, V.G., Carmeliet, P., and Kilpatrick, T.J. (2008). Gas6 deficiency increases oligodendrocyte loss and microglial activation in response to cuprizone-induced demyelination. J Neurosci 28, 5195– 5206. 10.1523/JNEUROSCI.1180-08.2008.

55. Nugent, A.A., Lin, K., van Lengerich, B., Lianoglou, S., Przybyla, L., Davis, S.S., Llapashtica, C., Wang, J., Kim, D.J., Xia, D., et al. (2020). TREM2 Regulates Microglial Cholesterol Metabolism upon Chronic Phagocytic Challenge. Neuron 105, 837–854.e9. 10.1016/j.neuron.2019.12.007.

56. Wang, N., Wang, M., Jeevaratnam, S., Rosenberg, C., Ikezu, T.C., Shue, F., Doss, S.V., Alnobani, A., Martens, Y.A., Wren, M., et al. (2022). Opposing effects of apoE2 and apoE4 on microglial activation and lipid metabolism in response to demyelination. Mol Neurodegener 17, 75. 10.1186/s13024-022-00577-1.

57. Ennerfelt, H., Frost, E.L., Shapiro, D.A., Holliday, C., Zengeler, K.E., Voithofer, G., Bolte, A.C., Lammert, C.R., Kulas, J.A., Ulland, T.K., et al. (2022). SYK coordinates neuroprotective microglial responses in neurodegenerative disease. Cell 185, 4135–4152.e22. 10.1016/j.cell.2022.09.030.

58. Deng, Y., Kim, B., He, X., Kim, S., Lu, C., Wang, H., Cho, S.-G., Hou, Y., Li, J., Zhao, X., et al. (2014). Direct visualization of membrane architecture of myelinating cells in transgenic mice expressing membrane-anchored EGFP. Genesis 52, 341–349.

59. Yona, S., Kim, K.-W., Wolf, Y., Mildner, A., Varol, D., Breker, M., Strauss-Ayali, D., Viukov, S., Guilliams, M., Misharin, A., et al. (2013). Fate mapping reveals origins and dynamics of monocytes and tissue macrophages under homeostasis. Immunity 38, 79–91. 10.1016/j.immuni.2012.12.001.

60. Lu, Q., Gore, M., Zhang, Q., Camenisch, T., Boast, S., Casagranda, F., Lai, C., Skinner, M.K., Klein, R., Matsushima, G.K., et al. (1999). Tyro-3 family receptors are essential regulators of mammalian spermatogenesis. Nature 398, 723–728. 10.1038/19554.

61. Madisen, L., Zwingman, T.A., Sunkin, S.M., Oh, S.W., Zariwala, H.A., Gu, H., Ng, L.L., Palmiter, R.D., Hawrylycz, M.J., Jones, A.R., et al. (2010). A robust and high-throughput Cre reporting and characterization system for the whole mouse brain. Nat Neurosci 13, 133–140. 10.1038/nn.2467.

